# Prebiotically Plausible ‘Patching’ of RNA Backbone Cleavage Through a 3′-5′ Pyrophosphate Linkage

**DOI:** 10.1101/731125

**Authors:** Tom H. Wright, Constantin Giurgiu, Aleksandar Radakovic, Derek K. O’Flaherty, Lijun Zhou, Jack W. Szostak

**Affiliations:** Howard Hughes Medical Institute, Department of Molecular Biology, and Center for Computational and Integrative Biology, Massachusetts General Hospital, 185 Cambridge Street, Boston, Massachusetts 02114, United States

## Abstract

Achieving multiple cycles of RNA replication within a model protocell would be a critical step towards demonstrating a path from prebiotic chemistry to cellular biology. Any model for early life based on an ‘RNA world’ must account for RNA strand cleavage and hydrolysis, which would degrade primitive genetic information and lead to an accumulation of truncated, phosphate-terminated strands. We show here that cleavage of the phosphodiester backbone is not an endpoint for RNA replication. Instead, 3′ -phosphate terminated RNA strands are able to participate in template-directed copying reactions with activated ribonucleotide monomers. These reactions form a pyrophosphate linkage, the stability of which we have characterized in the context of RNA copying chemistry. We found that the pyrophosphate bond is relatively stable within an RNA duplex and in the presence of chelated magnesium. Under these conditions, pyrophosphate-RNA can act as a temporary ‘patch’ to template the polymerization of canonical ribonucleotides, suggesting a plausible non-enzymatic pathway for the salvage and recovery of genetic information following strand cleavage.

## Introduction

The RNA world hypothesis proposes an early stage in the evolution of life in which RNA was responsible for both catalysis and genetic inheritance^1–4^. In existing biology, ribonucleotide monomers are joined to form polynucleotides in a template-directed process by the enzyme-catalyzed reaction of 3′-hydroxyl groups with nucleoside 5′-triphosphates, forming a 3′-5′ phosphodiester-linked backbone. Before the evolution of protein enzymes, a replicase made of RNA may have catalyzed RNA synthesis, a possibility explored by the *in vitro* selection and evolution of ribozymes with ligase^5^ and polymerase^6–8^ activity. A more challenging problem is how these ribozymes could have emerged from a non-enzymatic process of chemical RNA replication. Chemical RNA replication starts with the binding of energy-rich activated ribonucleotides to an RNA template through Watson-Crick base pairing. The ribonucleotide monomers react with each other to form phosphodiester bonds, leading to a double-stranded RNA duplex; subsequent amplification requires separation of the strands, so that they can act as templates for further cycles of copying. Mechanistic studies which revealed the critical role of 5′-5′ imidazolium-bridged dinucleotides as the active species in primer extension^9^, and the discovery of 2-aminoimidazole as a superior 5′-phosphate activating group^10,11^, have recently enabled the copying of short mixed sequence RNA templates within vesicle models of early cells^12^. Despite this progress, the chemical copying of RNA is not currently able to produce RNA products of the length and complexity necessary to sustain ribozyme evolution^13,14^.

Any model proposing RNA or RNA-like polymers as early carriers of genetic information must also account for cleavage of the phosphodiester backbone^15^, which competes with copying chemistry. The cleavage of RNA strands results in a mixture of 2′ and 3′ phosphate terminated strands via an initial transesterification reaction followed by hydrolysis of a cyclic phosphate intermediate (Figure 1a). Cleavage of RNA strands is greatly accelerated by divalent metal cations, such as magnesium, that are also required for the copying chemistry to proceed at a reasonable rate. This degradation pathway constrains plausible rates of non-enzymatic RNA synthesis and would result in a net loss of material, and thus information, from a primordial genetic system if there were no mechanism for recycling of the phosphate-terminated strands^15^. The Sutherland lab^16^ has demonstrated a prebiotically plausible cycle of backbone repair that converts 2′-5′ to 3′-5′ phosphodiester linkages via iterative cycles of strand cleavage and repair, in which a 3′ phosphate plays a central role as a substrate for ligation chemistry. The identification of complementary pathways for RNA salvage or repair following strand cleavage^16,17^, or the discovery of additional functions for phosphate-terminated strands^18^, could further strengthen the case for RNA as the earliest carrier of genetic information.

**Figure 1.**
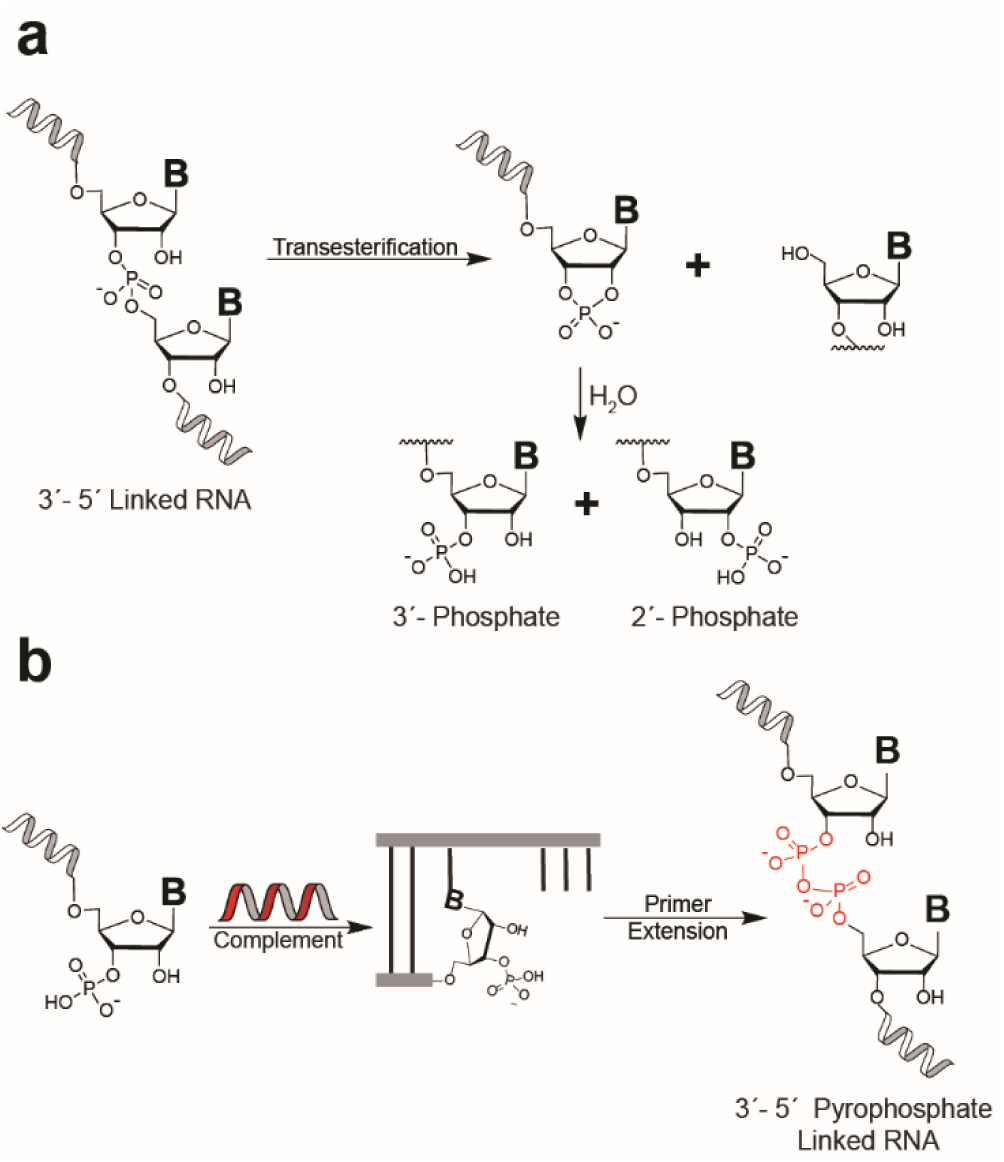
RNA strand cleavage and possible recovery of genetic information via pyrophosphate bond formation. (a) The cleavage of 3′-5′ phosphodiester-linked RNA leads to a mixture of 2′ and 3′ monophosphate terminated strands, via a cyclic phosphate intermediate. (b) Non-enzymatic primer extension starting from a 3′-monophosphate leads to 3′-5′ pyrophosphate-linked RNA, a possible mechanism for salvage of truncated RNA strands.

Consideration of potentially prebiotic nucleoside phosphorylation reactions provides further motivation for studying terminally phosphorylated RNA. Phosphorylation of nucleosides typically gives varying mixtures of 5′-, 3′- and 2′-phosphorylated products as well as the 2′:3′-cyclic phosphate. Although regioselective 5′-phosphorylation has been demonstrated, specific conditions such as the addition of borate minerals^19^ or the use of gas-phase reactions are required^20^. The diversity of pathways to nucleoside phosphorylation suggests that a mixture of phosphorylation states may have been likely at the monomer level. Heterogeneity of phosphorylation state at the 2′ and 3′ position of nucleotide monomers could therefore be incorporated into RNA strands via templated copying or non-templated reactions, in addition to the hydrolytic pathway.

Considering this implied presence of terminal phosphates in any ‘RNA world’ scenario and bearing in mind the greater nucleophilicity of phosphate relative to hydroxyl groups, we became interested in exploring whether phosphate-terminated RNA primers could participate in RNA copying chemistry. We hypothesized that the reaction of terminally phosphorylated primers with incoming nucleotides could lead to a mechanism for preservation of genetic information via formation of a pyrophosphate linkage (Figure 1b). Early work from Schwartz and Orgel demonstrated that 3′-phosphorylated, 2′-deoxy, imidazole-activated nucleotides efficiently polymerize on DNA templates to form 3′-5′ pyrophosphate-linked oligomers^21^. In turn, the 3′-5′ pyrophosphate-linked oligomers could be used as templates for further synthesis, again employing activated 3′-phosphate, 2′-deoxy monomers, indicating that a pyrophosphate-linked backbone may not preclude cycles of non-enzymatic replication^22^. Unfortunately, these early results were never extended to RNA-like systems.

Here, we report the results of our initial investigations into the behavior of phosphate-terminated RNA primers in non-enzymatic primer extension. We confirm that terminally phosphorylated RNA primers participate in primer extension reactions, leading to RNA polymers containing a pyrophosphate linkage. We have undertaken a thorough study of the stability of a single pyrophosphate linkage embedded within an RNA strand, which revealed pronounced lability towards cleavage reactions in the presence of magnesium ions. However, similarly to ‘native’ RNA, the pyrophosphate linkage is protected from strand cleavage in the context of a duplex and in the presence of magnesium chelators. These observations enabled us to survey the kinetic parameters of primer extension from phosphate-terminated primers and following incorporation of a pyrophosphate bond. We further demonstrate that pyrophosphate-linked RNA can function as a template to direct the polymerization of canonical ribonucleotides, pointing towards a possible non-enzymatic salvage pathway for primordial RNA.

## Results

We first synthesized RNA primers terminated with either a 2′ or 3′ monophosphate, using a solid phase approach (SI Figure 1a). Using strong-anion exchange chromatography^16^, we confirmed the regioisomeric purity of the individual primers; no contamination with the alternative terminal phosphate was observed in either case (SI Figure 1b). We therefore tested the 2′ and 3′ phosphorylated primers in a primer-extension assay (Figure 2), employing a templating region with the sequence 3′-CCCA-5′, which can be extended efficiently by addition of activated guanosine and uridine ribonucleotide monomers. Citrate-chelated magnesium was used in these assays as it enables RNA copying chemistry to proceed within vesicles composed of fatty acids^23^, which are the most likely candidates for protocell membranes but are destabilized by Mg^2+^ at low millimolar concentrations^24^. The two phosphorylated primers, and a non-phosphorylated control, were annealed separately with the template, incubated with 50 mM Mg^2+^, 200 mM citrate and 10 mM of both guanosine 2-aminophosphorimidazolide (2AImpG) and uridine-2-aminophos-phorimidazolide (2AImpU) and monitored over the course of 24 hours. All three primer-template duplexes were extended, although extension in the case of the 2′-phosphate terminated primer was very poor. Surprisingly, the 3′ phosphate-terminated primer gave comparable extension to that obtained in the 3′-hydroxyl system, with 81% of +3 products after 24 hours (cf. 87% of +3 product for 3′-hydroxyl).

**Figure 2.**
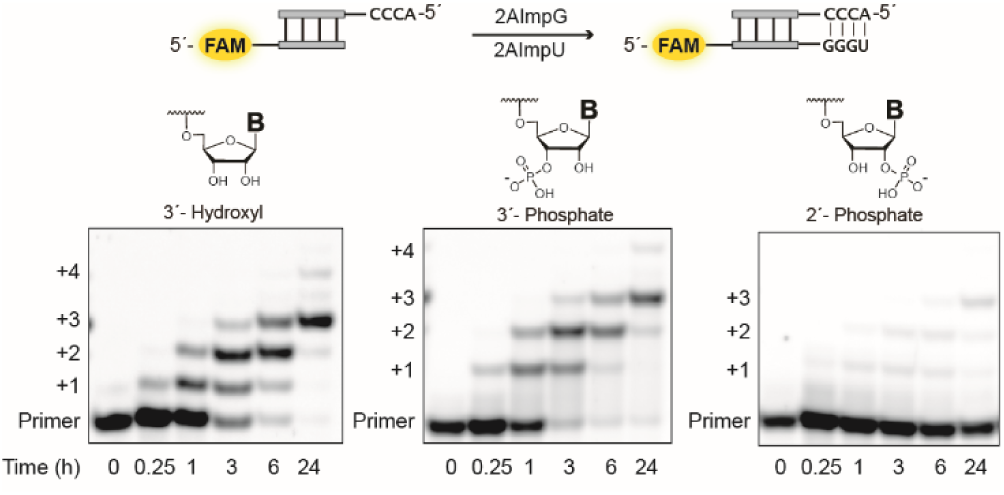
Efficient non-enzymatic copying of RNA commences from a primer with a terminal 3′-monophosphate. The time course of primer extension with guanosine 2-aminophosphorimidazolide (2AImpG) and uridine-2-aminophosphorimidazolide (2AImp-U) monomers was monitored using polyacrylamide gel electrophoresis (PAGE). All reactions were performed at pH 8.0, 200 mM HEPES, 50 mM Mg^2+^ and 200 mM citrate, with 10 mM each of guanosine 2-aminophosphorimidazolide and uridine-2-aminophosphorimidazolide.

We hypothesized that extension of the phosphate-terminated primers could be due to formation of a pyrophosphate linkage by reaction of the 2′ or 3′ terminal phosphate with the activated 5′-phosphate of the incoming monomer. However, we aimed to exclude an alternate possibility in which the reaction proceeds through the non-phosphorylated hydroxyl, as chemical primer extension can proceed via either the 2′ or 3′ hydroxyl group^25^. To confirm the presence of a pyrophosphate linkage in the reaction products derived from the 3′-phosphorylated primer, we made use of the fact that RNAse A is known to cleave pyrophosphate bonds^26^ while leaving 2′-5′ phosphodiester linkages intact^25^ (Figure 3a). The primer utilized was designed so that it is cleaved by RNAse A at a single site, after the first bond-forming step of the primer extension reaction. Thus, for the 3′-terminal phosphate reaction, we expected cleavage if the addition of the first monomer occurs through the 3′ phosphate and no cleavage if the reaction instead proceeded through the 2′ hydroxyl. Treating a primer extension reaction mixture with increasing concentrations of RNAse A resulted in cleavage of the +1 to +3 bands and the appearance of a band with the same gel mobility as the original primer. As RNAse A requires a free 2′-OH for cleavage and should lead to regeneration of the 3′-monophosphate, this result rules out formation of a 2′-5′-phosphodiester bond during primer extension and strongly suggests the presence of a 3′-5′-pyrophosphate linkage.

**Figure 3.**
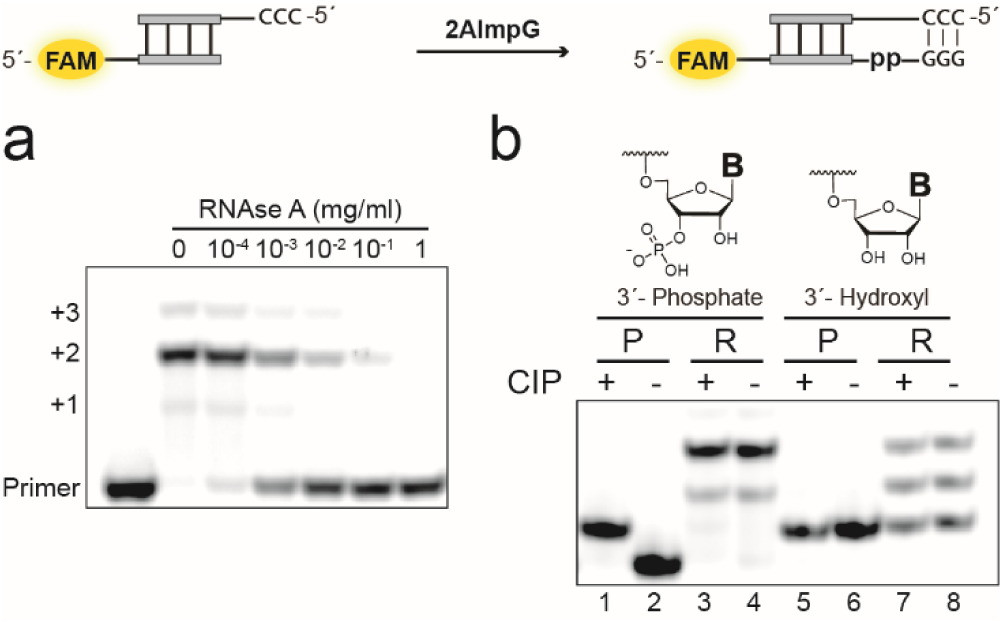
Confirmation of 3′-5′ pyrophosphate linkage by enzymatic digestion assays. (a) PAGE analysis of RNAse treatment. RNAse A treatment of a primer extension reaction commenced from a primer bearing a terminal 3′-monophosphate leads to regeneration of the primer. (b) PAGE analysis of calf intestinal phosphatase (CIP) treatment of primer extension reaction mixtures. ‘P’ refers to primer and ‘R’ refers to reaction mixture derived from primer extension. Lanes 1 and 2: 3′-phosphory-lated primer standard. Lanes 3 and 4: Reaction mixture from extension of the 3′-phosphorylated primer with (Lane 3) and without (Lane 4) CIP added. Lanes 5 and 6: Non-phosphorylated RNA primer. Lanes 7 and 8: Reaction mixture from extension of the same RNA primer.

If primer extension proceeded from the free terminal hydroxyl and not from the terminal phosphate group, a phosphate monoester internal to the RNA chain would remain in the +1 and higher extension products. To test this hypothesis, we treated the reaction mixtures with calf intestinal phosphatase (CIP), which has been shown to cleave internal phosphate monoesters^27^ (Figure 3b). For reaction mixtures derived from the 3′-phosphorylated primer, gel bands corresponding to extended products were not affected by CIP treatment. Taken together, these results obtained from enzymatic digestion support formation of a pyrophosphate linkage during primer extension initiated from a terminal phosphate and rule out the alternative possibility of a 2′-5′ phosphodiester-linked structure.

To further investigate the properties of pyrophosphate-linked RNA, we required a method to generate pure, single-stranded RNA containing site-specific pyrophosphate linkages, for use both as primers and as templates in downstream studies. Notably, there exist no synthetic or enzymatic routes to produce RNA containing defined pyrophosphate linkages. We thus devised a strategy that enabled us to generate pure (>95% by PAGE analysis) RNA strands containing a single pyrophosphate linkage (Figure 4). Our approach begins with a primer extension reaction using a 3′-phosphate terminated primer and a mixed sequence template which allows us to control the number and identity of added nucleotides simply by adjusting the reaction times or adding/removing nucleotide phosphorimidazolides from the reaction mixture. Critically, we use a DNA-RNA hybrid template in which the region to be copied is RNA but the remainder of the template is DNA, to allow for DNase digestion of the template which facilitates recovery of the modified primer. Incubation of the primer template duplexes with 2-aminoimidazole activated monomers leads to robust conversion of the primer to extended products in which the +1 nucleotide is connected to the RNA primer by a pyrophosphate linkage. Following desalting to remove unreacted monomer, a 5′-phosphorylated ligator RNA oligonucleotide is annealed and ligated to the pyrophosphate-containing primer by T4 RNI2 ligase. The ligation step combined with design of the primer and template sequence allows for the generation of essentially any RNA sequence in the final product. After ligation, treatment of the reaction mixture with DNase followed by gel purification affords reasonable yields (typically 20-30%, based on known input of primer) of high purity (>95% by PAGE analysis), single stranded ‘pyrophosphate-containing RNA’. To produce ssRNA species containing a terminal pyrophosphate (Figure 4), a template containing only a single binding site for the activated G-G dinucleotide intermediate is employed using a 5′-CC-3′ templating region and the ligation step is omitted. The +1 primer extension product, bearing a terminal pyrophosphate linkage, can be isolated directly following DNase digestion. Although a recent report detailed the synthesis of pyrophosphate-linked DNA via a solid-phase synthesis approach^28^, our method represents the first route to RNA containing defined pyrophosphate bonds.

**Figure 4.**
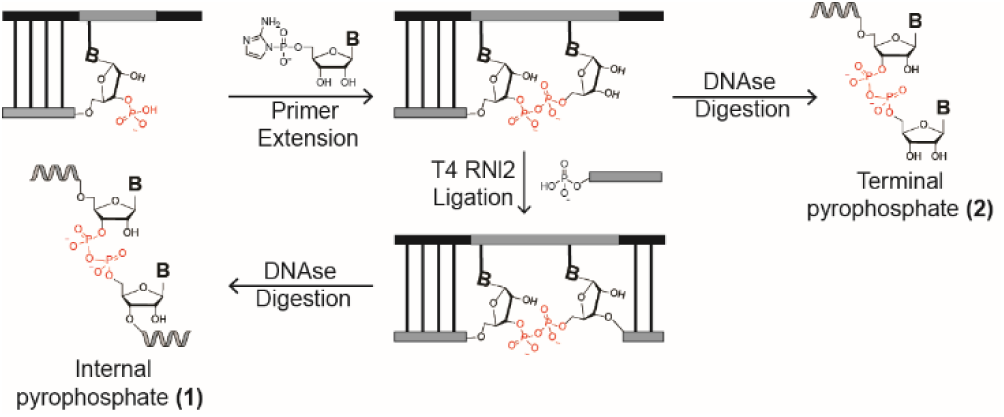
Synthetic strategy to access RNAs containing a single, defined pyrophosphate linkage. An RNA primer containing a terminal 3′-phosphate monoester is annealed to a chimeric DNA/RNA template. Primer extension with activated guanosine monomers generates a 3′-5′ pyrophosphate linkage. Digestion of the DNA residues of the template with Turbo DNase generates a ssRNA containing a terminal pyrophosphate linkage. Alternatively, T4 RNA ligase 2 can be used to ligate an RNA oligonucleotide downstream in excellent yields (>95%). The same DNase treatment can then be used to generate ssRNA containing an internal pyrophosphate linkage. Grey: RNA, Black: DNA. See the Supporting Information for detailed experimental procedures.

We were interested in exploring the relative stability of the pyrophosphate linkage within RNA, to determine whether it could support the chemical replication of genetic information, and whether it could in principle enable ribozyme function. We prepared two single-stranded RNAs containing either an internal (AAGGGAAGAAGC-pp-GGGCUAGCAUGAC **1**) or terminal pyrophosphate linkage (AAGGGAAGAAGC-pp-G **2**), using the strategies outlined above. The pyrophosphate-RNAs were then incubated at 22 °C in a pH 8.0 solution (conditions typical for primer extension reactions) in the presence or absence of magnesium, which was included as either the free cation or in citrate-chelated form (Table 1). We included citrate as it protects both RNA and fatty acid vesicles from magnesium-induced degradation, while still allowing RNA copying reactions to proceed, albeit at a reduced rate^23^. In the presence of 1 mM EDTA, which should chelate trace divalent cations, the pyrophosphate linkage was relatively stable, with a half-life of 16 days. In contrast, incubation with free magnesium (50 mM) led to rapid cleavage of the pyrophosphate bond, with a half-life measured to be on the order of minutes for both internal and terminal pyrophosphate linkages. Even at low millimolar concentrations of magnesium ions, the half-life of the pyrophosphate bond was only extended slightly (SI Table 1). The product of strand cleavage was determined by LC-MS (SI Figure 2a) and CIP digestion (SI Figure 2b) to be the 2′-3′-linked cyclic phosphate, consistent with a mechanism in which the α-phosphorus atom is subject to nucleophilic attack by the adjacent 2′-hydroxyl group. Incubating the pyrophosphate-RNA with magnesium citrate afforded modest protection, extending the half-life of the linkage from minutes to hours. Notably, the reduction in the rate of cleavage afforded by citrate chelation is much larger than the effect of chelation on the rate of the copying chemistry^23^.

**Table 1.** Half-life of 3′-5′ pyrophosphate linked RNA*

|  | ssRNA | dsRNA | Stabilization |
| --- | --- | --- | --- |
| *Internal pyrophosphate* |  |  |  |
| Mg<sup>2+</sup> | 0.12(1) | 690(20) | 5500(200) |
| Mg-citrate | 3.7(1) |  |  |
| Citrate | 30 (1) |  |  |
| EDTA | 390 (20) |  |  |
| *Terminal pyrophosphate* |  |  |  |
| Mg<sup>2+</sup> | 0.20 (1) | 26(1) | 130 (10) |
| Mg-citrate | 3.7(2) |  |  |

\*The half-life values are presented with the standard error of the mean and reported in hours.

Duplex formation has been shown to protect 3′-5′ linked phosphodiester bonds in RNA from strand cleavage, while promoting cleavage in the case of 2′-5′ bonds^29,30^. This observation has been used to explain the ultimate ‘selection’ of the canonical 3′-5′ linkage in extant RNA. We therefore examined the effect of duplex formation on the rate of cleavage of the pyrophosphate linkage, to determine whether pyrophosphate bonds, once formed, could be retained in RNA duplex structures and ultimately serve as components of templates for further copying cycles. For the RNA 25mer **1** containing an internal pyrophosphate, addition of the complementary strand ‘buries’ the pyrophosphate bond in an extensive duplex region. Incubating this duplex with free Mg^2+^ revealed a significant stabilization effect, with the half-life of the pyrophosphate bond increased from minutes to 28 days; a stabilization factor of 5500. Protection in the case of the terminal pyrophosphate was far more modest, consistent with reduced helicity at the primer terminus.

Together, our degradation studies constrain the conditions under which the pyrophosphate linkage could support the storage and propagation of early genetic information. Pyrophosphate retention is possible within a duplex but is strongly disfavored in single-stranded RNA in the presence of Mg^2+^. The fate of a pyrophosphate bond at the terminal end of a duplex will depend on the relative rates of downstream extension or ligation reactions and strand cleavage and is thus expected to also depend on the RNA sequence (see Discussion below).

A critical factor determining the proportion of pyrophosphate-containing RNAs within a prebiotic population of polynucleotides would be the relative rates of reaction of phosphate and hydroxyl terminated primers with activated nucleotides. We were therefore interested in comparing the kinetics of primer extension for the initial reaction step. We measured the rates of primer extension on a template designed to provide a single binding site for C*C dimer (5′-5′ aminoimidazolium-bridged cytidine dinucleotide), the reactive species in primer extension using 2-aminoimidazole activated cytidine ribonucleotides (Figure 5). Using varying concentrations of C*C, we obtained Michaelis-Menten parameters for reactions using primers terminated in either a hydroxyl or monophosphate group at the 3′ position. We performed these experiments with magnesium-citrate, which protects against pyrophosphate degradation on the time-scales examined. Notably, we observed both higher v_max_ (14.1 h^−1^ vs. 4.4 h^−1^ obtained for hydroxyl) and K_M_ values (62.6 mM vs. 8.4 mM for hydroxyl) for the phosphate-terminated primer (Figure 5). The observed binding defect could be due to either charge repulsion between the negatively charged phosphate and the dimer species or to steric hindrance. The rate enhancement may be rationalized by consideration of the active nucleophile in each case. The actual nucleophilic species in primer extension with ‘native’ RNA is likely a Mg-bound alkoxide, which is poorly populated at pH 8^13^. The nucleophile for the 3’-phosphorylated primer is most likely a phosphate monoester dianion, which is more highly populated at pH 8^31^. The difference in the population of these two nucleophiles under primer extension conditions may therefore explain the observed rate difference.

**Figure 5.**
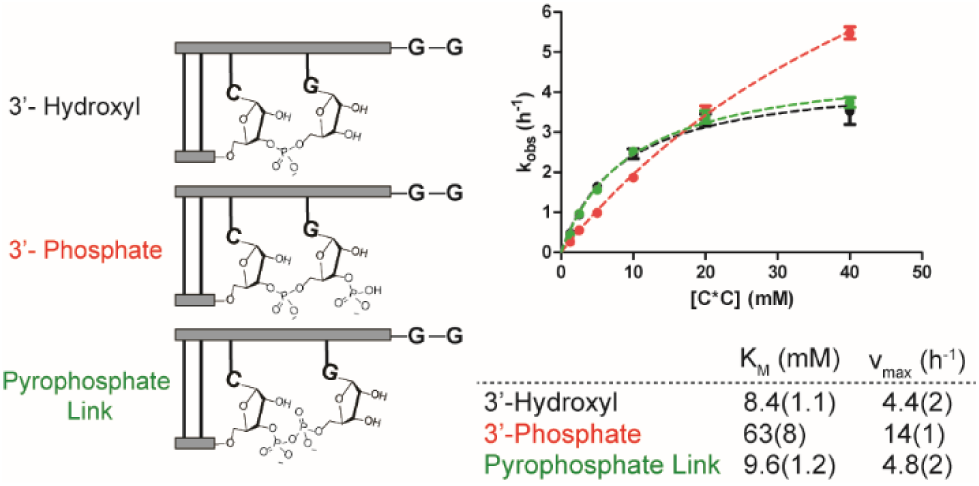
Michaelis-Menten analysis of non-enzymatic copying from 3′-hydroxyl and 3′-monophosphate terminated primers, and for a 3′-hydroxylated terminus following a pyrophosphate bond. Left: Schematic representation of the primer-template duplexes analyzed. Right: Michaelis-Menten curves and obtained kinetic parameters. Plotted is k_obs_ (h-^1^) against the concentration of C*C dimer. All reactions were performed at pH 8.0, 200 mM HEPES, 50 mM Mg^2+^ and 200 mM citrate.

Previous reports have demonstrated a stalling effect on primer extension following the incorporation of mismatched bases^32^ and certain non-canonical nucleotides^33^. Our examination of the stability of the +1 extended product indicates that downstream extension is required to afford protection against cleavage of the newly formed pyrophosphate bond. We therefore sought to quantify the relative rates of reaction for primers in which the terminal 3′ nucleotide is joined by either a pyrophosphate or phosphodiester bond (Figure 5). The kinetic parameters for incorporation of the next nucleotide were almost identical for the pyrophosphate and phosphodiester linked systems (v_max_ of 4.8 h^−1^ following a pyrophosphate linkage vs. 4.4 h^−1^ obtained for phosphodiester-linked). Importantly, the defect observed for binding of the imidazolium bridged dimer intermediate C*C to the 3′ phosphate terminated primer was almost completely ameliorated (K_M_ 9.6 mM vs. 8.4 mM for phosphodiester-linked). This result implies that after the initial pyrophosphate linkage formation, the downstream steps in primer extension proceed essentially as if the primer contains only native RNA.

If pyrophosphate-linked RNA can act as a template for primer extension with canonical, 3′-hydroxyl containing ribonucleotide monomers, a mechanism for re-enrichment of phosphodiester-linked RNA could operate over cycles of replication, as the altered backbone structure will not be passed on to daughter strands. We were therefore curious whether a single 3′-5′ pyrophosphate linkage internal to an RNA template influences the ability of the template to direct monomer incorporation. For these experiments, we prepared 20mer ssRNA **3** (AUCGAAGGGppGGCAACACGAC), which contains a single pyrophosphate linkage, as the template strand. The template was designed to contain a stretch of guanosine residues 5′-GppGGG-3′ such that we could directly compare the kinetic parameters of template-copying across different systems using the same C*C dimer employed in our previous kinetic experiments. We examined three cases that differ only in the position of the primer strand relative to the pyrophosphate bond within the template (Figure 6). In the first case, the pyrophosphate bond joins the two nucleotides of the dimer binding site (Figure 6a). In the second case, the pyrophosphate bond is located after the primer annealing site, such that the expected distance between the primer 3′-hydroxyl and the 5′-phosphate of the incoming dimer is greater than in the case of phosphodiester-linked RNA (Figure 6b). Finally, in the third case examined the primer is ‘clamped’ over the pyrophosphate linkage, rendering it internal to the primer-template duplex structure (Figure 6c). Using varying concentrations of C*C, we evaluated initial rates of primer extension and determined Michaelis-Menten parameters for the three cases using hydroxyl-terminated primers and either the pyrophosphate-containing template or a fully RNA template of the same sequence. For all three situations, extension could be observed, and rates determined, for concentrations of imidazolium bridged intermediate as low as 2 mM. This result indicates that pyrophosphate-linked RNA can indeed act as template for the incorporation of canonical ribonucleotide monomers (Figure 6). Binding of the imidazolium-bridged C*C dimer to the template was only strongly affected when the pyrophosphate linkage directly connects the two bases of the binding site (~7-fold increase in K_M_ observed, Figure 6a). However, in all three cases, the maximum rate (v_max_) values obtained were lower for the pyrophosphate-linked templates. When comparing the three cases, the observed rates of copying across the pyrophosphate template were higher, and the difference in maximum rate between RNA and pyrophosphate templates was less pronounced, when the primer ‘clamps’ over the pyrophosphate linkage (Figure 6c). This may imply that conformational distortion in the ternary complex of primer: template: bound dimer due to the pyrophosphate bond becomes less significant as the linkage is buried within an RNA duplex.

**Figure 6.**
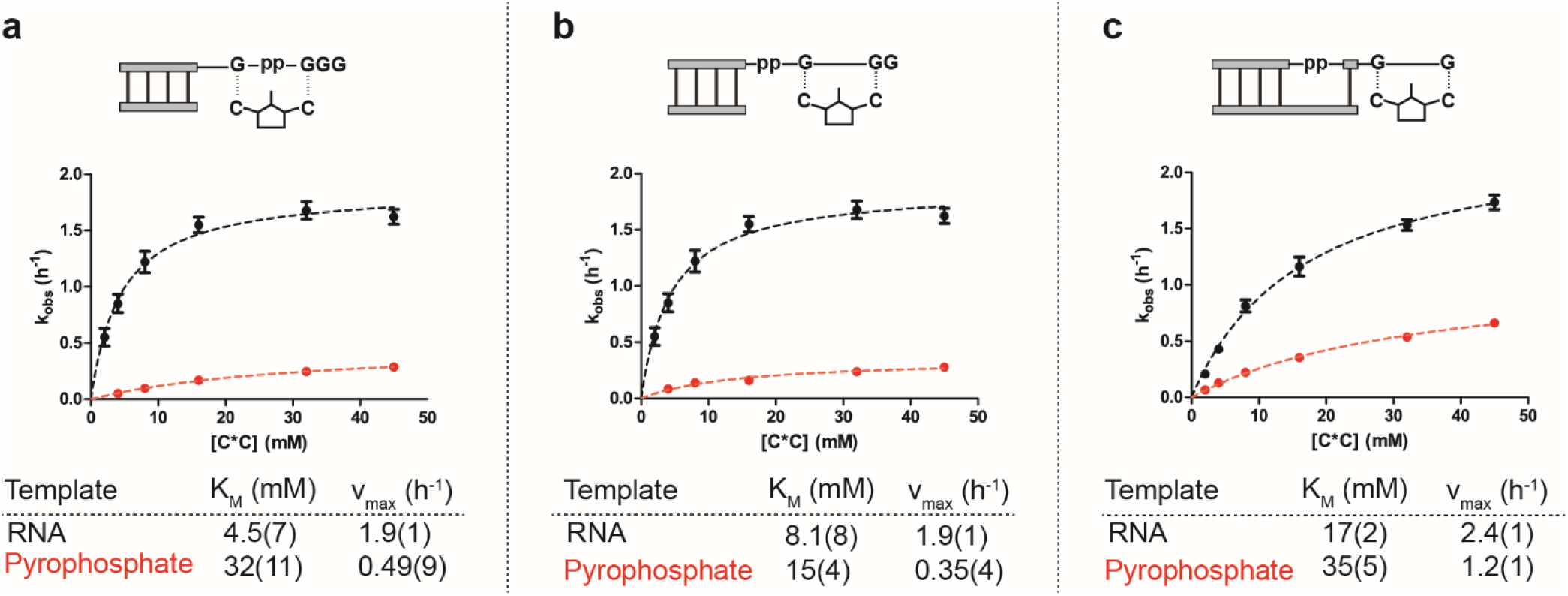
(a-c) Michaelis-Menten analysis of non-enzymatic copying across 3′-5′ pyrophosphate-linked template 3, compared with phosphodiester-linked control. Top: Schematic representation of the primer-template duplexes analyzed, showing the binding site for the C*C dimer in each case. Bottom: Michaelis-Menten curves and obtained kinetic parameters. Plotted is k_obs_ (h^−1^) against the concentration of C*C dimer. All reactions were performed at pH 8.0, 200 mM HEPES, 50 mM Mg2+ and 200 mM citrate. Further experimental details and data analysis procedures can be found in the Supporting Information.

## Discussion

We have found that RNA strand cleavage and hydrolysis is not a dead-end for primitive, RNA-based genetic systems. Instead, the 3′phosphorylated RNAs that result are able to participate in non-enzymatic primer extension when supplied with activated nucleotides. Upon reaction of phosphate-terminated primers with incoming nucleotides, a 3′-5′ pyrophosphate linkage is formed.

By surveying the kinetic parameters for both the synthesis and degradation of this linkage, we have been able to place reasonable constraints on the likelihood of pyrophosphate-linked RNA playing a role in early systems of genetic inheritance.

The kinetic stability of the phosphodiester linkage is a critical feature enabling the storage and use of genetic information^34^. In an RNA world dependent on ribozymes for catalysis, the rate of cleavage of the phosphodiester bond should also determine the lifetime, and thus biosynthesis requirements, of the functional machinery of the protocell^15^.

In the transition from prebiotic chemistry to the first genetic polymers, the phosphodiester linkage may have been ‘selected’ from a range of alternative backbones that were feasible from the standpoint of chemical reactivity but were not stable enough to support the long-term storage and transmission of genetic information or the functioning of ribozymes. Considering only canonical ribonucleotide monomers as precursors to primitive RNA oligonucleotides, four backbone linkages are possible which are derived from either non-enzymatic copying reactions or the cleavage pathways discussed in this manuscript: 3′-5′ and 2′-5′ phosphodiester linkages and the 3′-5′ and the 2′-5′ pyrophosphate linkages explored here. Our initial primer extension results suggest that 2′-5′ pyrophosphate linkages were unlikely to play a significant role in early forms of RNA-based genetics, as initiation of copying chemistry from a 2′-phosphate is strongly disfavored. In contrast, copying from 3′-phosphate terminated primers is robust and comparable to extension from 3′-hydroxyl groups.

When considering degradation reactions, the rate of cleavage of a 3′-5′ pyrophosphate linkage within single-stranded RNA in the presence of magnesium ions is orders of magnitude greater than that of either a 2′-5′ or 3′-5′ phosphodiester linkage^29,30^. However, duplex formation protects 3′-5′ phosphodiester-linked bonds from cleavage, by constraining RNA in a helical conformation that disfavors attack on the phosphodiester unit by the adjacent 2′-hydroxyl. We observed a similar effect for 3′-5′ pyrophosphate linkages, suggesting a similar mechanism of protection. In the case of 2′-5′ phosphodiester linkages, duplex formation has the opposite effect, promoting attack by the 3′-hydroxyl and leading to more rapid cleavage. We expect a similar effect to operate for 2′-5′-linked pyrophosphate bonds, although the inefficiency of primer extension from 2′-phosphates has prevented us from directly testing this hypothesis. The observation of enhanced stability for the 3′-5′ phosphodiester linkage in duplex structures has previously been used to support the idea of ‘selection’ over time for the more stable 3′-5′ linkage in the context of genetic inheritance^29^. Our results suggest that susceptibility to strand cleavage in the presence of magnesium ions, which are required for RNA folding and catalysis, could explain why biology does not currently employ 3′-5′ pyrophosphate linkages in the RNA backbone. However, before the emergence of enzymatic mechanisms for synthesis and proof-reading of genetic polymers, pyrophosphate linkages could have been tolerated as part of duplex structures and thus have contributed to the transmission of primitive genetic information.

We have shown that pyrophosphate linkages in template strands do not block template-directed primer extension. This result suggests pyrophosphate linkages may have been tolerated in primordial RNA-based genetics and raises the prospect of a non-enzymatic ‘salvage’ and proof-reading cycle for cleaved RNA (Figure 7). Following a cleavage(which could take place in a single strand or within a duplex structure), a cyclic phosphate terminus results and is further hydrolyzed to a mixture of 2′ and 3′ terminal phosphates, with the latter being slightly favored^35^. If a 3′-phosphorylated RNA is bound by a template strand, reaction with activated monomers could ‘patch’ the cleaved strand via formation of a pyrophosphate linkage. This would result in a duplex structure bearing a single pyrophosphate linkage, the lifetime of which depends on the relative rate of pyrophosphate cleavage compared to primer extension or ligation reactions. Cleaved pyrophosphate bonds can simply re-enter the cycle while those sequences able to extend efficiently (or undergo rapid ligation reactions) will ‘bury’ the labile pyrophosphate bond within a duplex structure, affording some protection from cleavage. For regeneration of the original phosphodiester-linked RNA strand, strand separation and copying of the pyrophosphate-linked RNA (now acting as template) is required. If only canonical ribonucleotide monomers are available, we have demonstrated that copying over pyrophosphate linkages is feasible, albeit slower than for purely phosphodiester-linked RNA. Non-enzymatic separation of RNA strands and multiple cycles of copying (replication) have yet to be demonstrated experimentally^14,36^. Assuming such a cycle, phosphodiester bond formation over a pyrophosphate template would re-enrich canonical RNA in a crude, non-enzymatic form of backbone proofreading. The extreme susceptibility of pyrophosphate linkages to metal-ion catalyzed cleavage in single-strands renders this pathway unlikely under a model of non-enzymatic replication that requires temperature cycling, as this promotes a fully single-stranded state under conditions likely to promote cleavage. However, if a strand displacement synthesis is possible, temperature cycling is not required, and the salvage pathway may be feasible.

**Figure 7.**
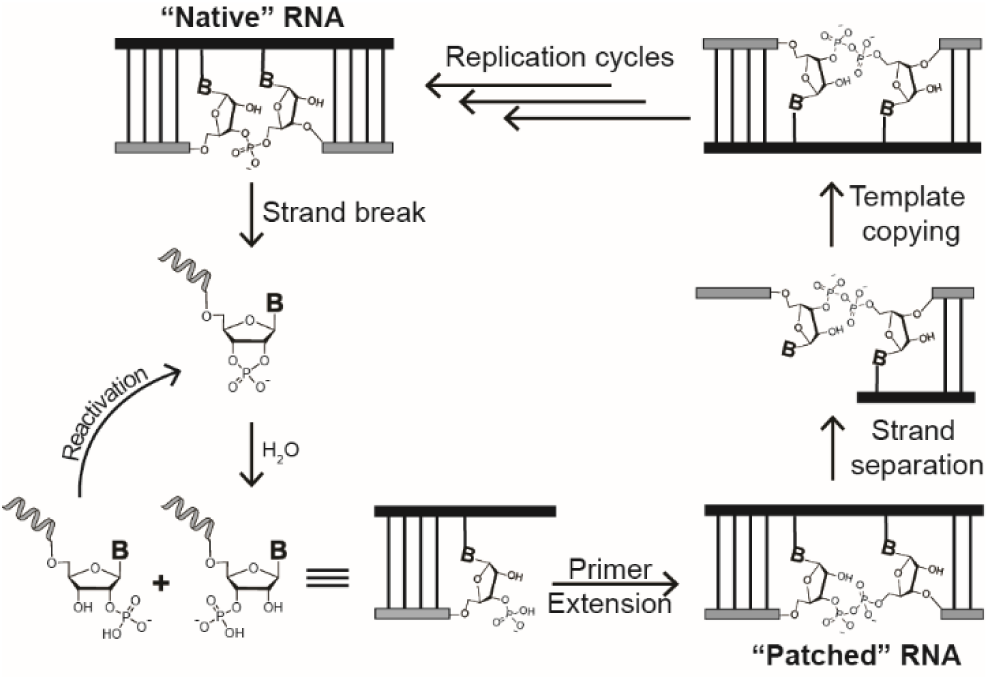
A non-enzymatic pathway for the recovery of genetic information via a pyrophosphate ‘patch’. (From top left) Following a cleavage event, a cyclic phosphate terminus is formed which hydrolyzes to a mixture of 3′ and 2′ monophosphate-terminated strands. Primer extension from a 3′-monophosphate terminus leads to duplex RNA containing a single 3′-5′ pyrophosphate linkage. Strand displacement and templated copying with canonical ribonucleotides leads to re-enrichment of ‘native’, 3′-5′ phosphodiester-linked RNA over multiple cycles of replication. The unreactive 2′ monophosphate-terminated strands can be reactivated to the cyclic phosphate and re-enter the cycle.

Mariani et al.^16^ have demonstrated a different cycle of backbone repair that converts 2′-5′ to 3′-5′ phosphodiester linkages via iterative cycles of strand cleavage, 2′-acetylation and non-enzymatic RNA ligation^17^.

Such a cycle provides a pathway for the repair of genetic information that also relies on the formation of a 3′-phosphate as the critical intermediate. In conditions that promote the chemical activation of phosphate groups, ligation with a 5′-hydroxyl is thus a complementary pathway to that described in this manuscript.

Under such activating conditions, if a 5′-phosphate is present on a bound ligator, pyrophosphate formation should also occur. Hydrolysis of a 3′-5′ pyrophosphate linkage produces such a 5′-phosphate on the downstream cleaved fragment, which could feed back into the system as an input for repeated ligation with a 3′-phosphate or ligation to an oligo bearing a 3′-hydroxyl. A network of interconnecting salvage and ‘repair’ pathways for RNA may therefore operate under non-enzymatic conditions, all of which depend on the unique reactivity of 3′-phosphates generated upon hydrolytic cleavage.

Acetylation of the vicinal 2′-hydroxyl group of a terminal 3′-phosphate before primer extension^16,17^ should render the resultant pyrophosphate linkage stable to strand cleavage as the 2′-hydroxyl would be incapable of transesterification. Such prebiotically plausible ‘protection’ could greatly enhance the retention of pyrophosphate linkages within a population of RNAs and bolster the chances of repair by enhancing the kinetic stability of pyrophosphate-containing templates. As a proof of principle that protecting the 2′-hydroxyl would not inhibit primer extension from a terminal 3′-monophosphate, while reducing the problem of strand cleavage, we synthesized a primer with a 2′-OMe, 3′-phosphorylated terminus and compared its activity with our original 2′-OH, 3′-phosphorylated primer in a primer extension assay with free magnesium cations (SI Figure 3). Over 18 hours, both primers were extended up to +6 products, with a slightly greater yield of +6 products observed for the 2′-O-methylated primer. Pleasingly, minimal remaining primer (5%) was observed for the 2′-O-methylated system during the course of the assay. In contrast, the reaction with a free 2′-hydroxyl yielded 22% of a product that co-migrates with the original primer band, consistent with cleavage leading to a cyclic phosphate terminated strand (evidence for the co-migration of our 3′-phosphorylated primer and its derived cyclic phosphate is presented in SI Figure 2b). The fraction of remaining primer observed for the 2′-O-methylated system correlates with that observed in identical reactions conducted with nonphosphorylated RNA, in which native phosphodiester bonds are formed that do not appreciably cleave on this time scale. This result suggests that prebiotically-plausible 2′-OH protection could indeed facilitate the increased retention of pyrophosphate bonds in early RNA polymers and perhaps expand potential roles for such linkages in the RNA world. Work towards this end is currently underway in our laboratory.

This study only examined primer extension using canonical ribonucleotides reacting with phosphate groups introduced to the 3′ ends of primers by solid-phase synthesis. A remaining question is the efficiency of primer extension using 2′ or 3′-phosphorylated monomers. Schwartz and Orgel have shown that 3′-monophosphate, 2′-deoxy modified monomers can polymerize on DNA templates to form 3′-5′ pyrophosphate-linked oligomers^21^. Isolated pyrophosphate-linked oligonucleotides could also be used as templates for further polymerization, employing the same activated monomers^22^. These results imply that a full cycle of non-enzymatic replication may be possible in a system composed of phosphorylated monomers and pyrophosphate-linked genetic polymers. Here, we examined the consequences of a single pyrophosphate linkage in the context of RNA copying. Primer extension using 3′-phosphorylated ribonucleotide monomers, or in a system containing monomers with a mixture of phosphorylation states, should introduce multiple pyrophosphate linkages. The implications of multiple pyrophosphate bonds for the function of RNA, as both genetic material and catalyst, remain to be explored.

## ASSOCIATED CONTENT

### Supporting Information

Methods and materials, Figures S1-S3, and Table S1.

### Notes

The authors declare no competing financial interests.

## ACKNOWLEDGMENT

J.W.S is an investigator of the Howard Hughes Medical Institute. The authors would like to thank Dr. Li Li for helpful discussions. This work was supported in part by a grant (290363FY18) from the Simons Foundation to J.W.S., a grant from the NSF (CHE-1607034) to J.W.S. and a grant from NASA (NNX15AL18G) to J.W.S. D.K.O. is a recipient of a Postdoctoral Research Scholarship (B3) from the Fonds de Recherche du Quebec−Nature et Technologies (FRQNT), Quebec, Canada, and a Postdoctoral Fellowship from Canadian Institutes of Health Research (CIHR) from Canada.

