## Supplemental Information for "Prebiotically Plausible ‘Patching’ of RNA Backbone Cleavage Through a 3′-5′ Pyrophosphate Linkage"

CCIB 7215, Simches Research Center

185 Cambridge Street

Massachusetts General Hospital

Boston, Massachusetts 02114

### 1. Methods and Materials

#### 1.1 General Methods

All chemicals and yeast tRNA were purchased from Sigma-Aldrich (St. Louis, MO) unless otherwise noted. RNase A and TurboDNase were purchased from Thermo Scientific (Waltham, MA) and CIP and T4 RNl2 were purchased from New England Biolabs (Ipswitch, MA). All mononucleotide 5′-phosphorimidazolides, as well as the imidazolium-bridged dinucleotide intermediate, were synthesized in-house as previously reported[1,2].

#### 1.2 Oligonucleotide Synthesis and Characterization

All oligonucleotides used in this study were purchased from Integrated DNA Technologies (Coralville, IA) or prepared by solid-phase synthesis using an Expedite 8909 DNA/RNA synthesizer. Synthesizer reagents and phosphoramidites were purchased from Glen Research (Sterling, VA) and Chemgenes (Wilmington, MA). In-house prepared oligonucleotides were deprotected and purified by gel electrophoresis or by DMT-ON reverse phase silica column chromatography. The oligonucleotides were analyzed by high resolution mass spectrometry (HRMS) on an Agilent 6520 QTOF LC-MS. The regioisomeric purity of oligonucleotides containing terminal phosphate monoesters was determined by strong anion exchange high-pressure liquid chromatography. A Dionex DNAPac PA-100, 4 × 250 mm analytical column was used, at a flow rate of 1 mL/min., Flow rate: 1 mL/min. Buffer A was a solution of 10 mM sodium phosphate pH 11.5 and buffer B was a solution of 10 mM sodium phosphate pH 11.5 and 1 M NaCl. The oligonucleotides were detected at 260 nM.

#### 1.3 Primer extension reactions

The primer extension reactions with activated phosphorimidazolides in the presence of unchelated $Mg^{2+}$ were carried out as previously reported[3]. The reactions using Mg-citrate and activated phosphorimidazolides were carried out as reported in our previous study[4]. All reactions were performed in triplicate.

The Michaelis-Menten experiments were performed using the same conditions as the Mg-citrate primer extensions, but with purified imidazolium-bridged dimer instead of activated monoribonucleotides. Four timepoints were used to determine the rate at each concentration of imidazolium-bridged dimer. Each rate was calculated from three independent replicates, and the values were used to fit the Michaelis-Menten curve. The values presented in Figures 5 and 6 are the mean values of $K_M$ and $v_{max}$, reported with the standard error of the mean.

#### 1.4 Synthesis of pyrophosphate-linked RNA

**Primer extension**

The first step in the synthesis of pyrophosphate-linked RNA is carrying out a primer extension reaction using a 3′-phosphate terminated primer and a chimeric DNA/RNA oligonucleotide as a template. The primer extension reactions were performed at pH 8.0, 200 mM HEPES, 50 mM $Mg^{2+}$ and a primer concentration between 15-50 uM. The chimeric template was used in 1.5-fold excess to the primer. The reactions were run with 2AImpG concentrations between 20-40 mM for one hour at room temperature, followed by quenching with a solution of 0.5 M EDTA pH 8.0. The final amount of EDTA was 1.1x that of $Mg^{2+}$, to ensure that all of the magnesium was chelated in order to protect the pyrophosphate bond.

**Buffer exchange**

The quenched reaction was brought to a volume of 500 µL using nuclease-free water, after which the mixture was concentrated to a volume of approximately 50 µL using Amicon Ultra-0.5 mL 3K centrifugal filters, according to the supplier's instructions. The volume of the mixture was brought back up to 500 µL using nuclease-free water and then concentrated back to 50 µL. This step was repeated twice.

**T4 RNl2 Ligation**

The mixture was brought to a final concentration of 10 mM Tris pH 7.5, 0.2 mM EDTA and 2 equivalents of RNA ligator (relative to the primer). The reaction mix was annealed by heating at 70 °C for 3 minutes and then cooled to 12 °C at a rate of 0.1 °C/s. The mixture was diluted 10-fold in a solution containing 1x T4 RNl2 buffer and 1 U/µL T4 RNl2. The reaction was incubated at 37 °C until all of the extended primer was converted to the ligated product. We found that the incubation time depended on the position of the pyrophosphate linkage, such that if two nucleotides were added at the end of the primer only one hour of reaction time was required. However, if only one nucleotide had been added, the T4 ligation reaction needed 48 hours to go to completion.

**DNase Digestion and Gel Purification**

After the reaction was completed, 0.42 V of water, 0.067 V of TurboDNase 10x buffer and 0.1833 V of TurboDNase (2 U/µL) were added, and the mixture was incubated for an additional 15 minutes at 37 °C. The RNA was precipitated by adding 0.1 V of 3 M NaOAc pH 5.5 and 2.5 V of EtOH and incubated on dry ice for 30 minutes. The RNA pellet was collected by centrifugation at 14000 g and 4 °C for 30 minutes. The pellet was washed with cold 70% EtOH twice, and then resolubilized in 10 mM EDTA in 95% formamide (v/v). The pyrophosphate-linked RNA was purified by preparative polyacrylamide gel electrophoresis. A 20% gel was run at 25W at 4 °C for 2 hours, then the desired band was excised and eluted overnight at 4 °C in a 20 mM MES pH 6.0, 5 mM EDTA and 250 mM NaCl solution. The RNA was then precipitated again and resuspended in a 1 mM EDTA, 5 mM MES pH 6.0 solution and used in further experiments.

If the terminal pyrophosphate product is desired, the DNase digestion step and gel purification is carried out immediately after the buffer-exchange step.

#### 1.5 Stability of the pyrophosphate linkage

For the single-stranded RNA experiments the previously synthesized oligomer (1) was incubated at room temperature in 50 mM HEPES pH 8.0, 50 mM NaCl and either one of the following conditions: 1 mM EDTA, 50 mM $Mg^{2+}$, 200 mM citrate or 50 mM $Mg^{2+}$ and 200 mM citrate. The reactions were quenched at different timepoints by taking a 2 uL aliquot and adding it to 18 uL of 10 mM EDTA in 95% (v/v) formamide. The percentage of remaining (not cleaved) oligomer was determined by PAGE and the values were fitted to a first-order exponential decay to obtain the rate constant for the first-order cleavage reactions.

For the double-stranded RNA experiments, the oligomer (1) was first annealed with three equivalents of its reverse complement in a solution containing 50 mM Tris pH 8.0, 1 mM EDTA and 50 mM NaCl, by heating the mixture at 70 °C for 3 minutes, followed by cooling to 12 °C at a rate of 0.1 °C/s. The oligomer solution was then diluted 13.33-fold in the same conditions used for the single-stranded RNA experiments, and the rate constant determination process was identical.

Each reaction constant was calculated from two independent replicates and the mean half-life values are reported together with the standard deviation of the mean.

#### 1.6 RNase A assay

The RNase A assay was carried out as previously reported[3].

#### 1.7 CIP assay

A 20 μL aliquot of the primer extension reaction mixture (3 μM primer concentration) was mixed with 60 μL of ethanol and 4 μL of 0.5 M EDTA in a siliconized microcentrifuge vial. The vial was placed in a -80 °C freezer for 2 hours, after which it was centrifuged at 4 °C and 21,000 g for 30 minutes. The resulting pellet was washed twice with 60 μL 70% ethanol, and then dried under vacuum for 20 minutes. The pellet was resolubilized in 20 μL 50 mM Tris pH 7.5. A 3 μL aliquot was diluted to 20 μL in 1x CutSmart buffer and 1U/μL CIP. The reactions were incubated at 37 °C for 90 minutes, after which they were analyzed by polyacrylamide gel electrophoresis.

#### 2. Supplementary Figures

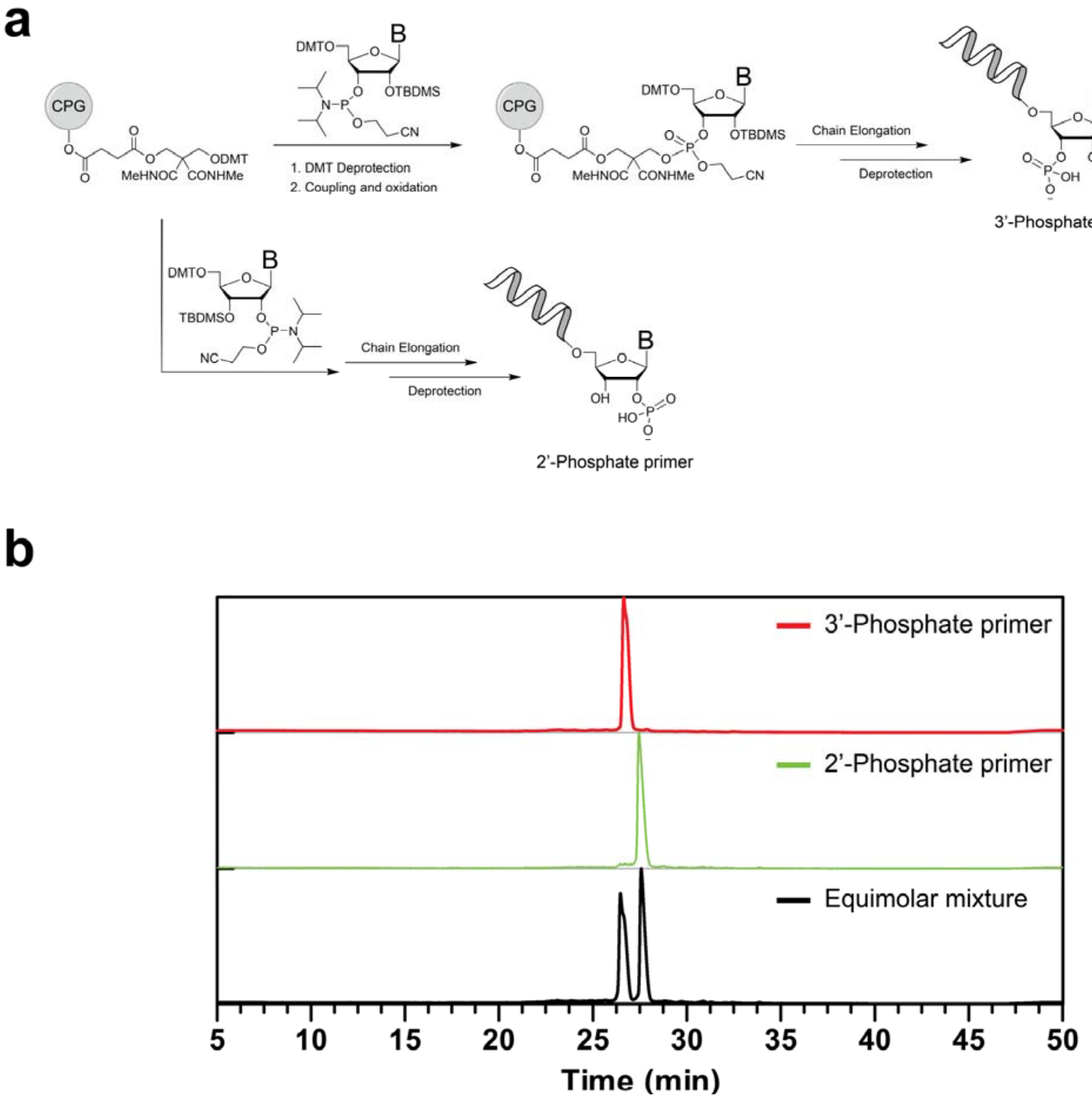

**Figure S1**. Synthesis and characterization of terminally phosphorylated primers. (a.) Synthetic scheme towards 3′ and 2′ monophosphate-terminated RNA strands via solid-phase synthesis. (b) Strong-anion exchange chromatography (SAX) was used to characterize the regioisomeric purity of the synthesized primers; no contamination with the alternative phosphoryl-regioisomer was observed in either case.

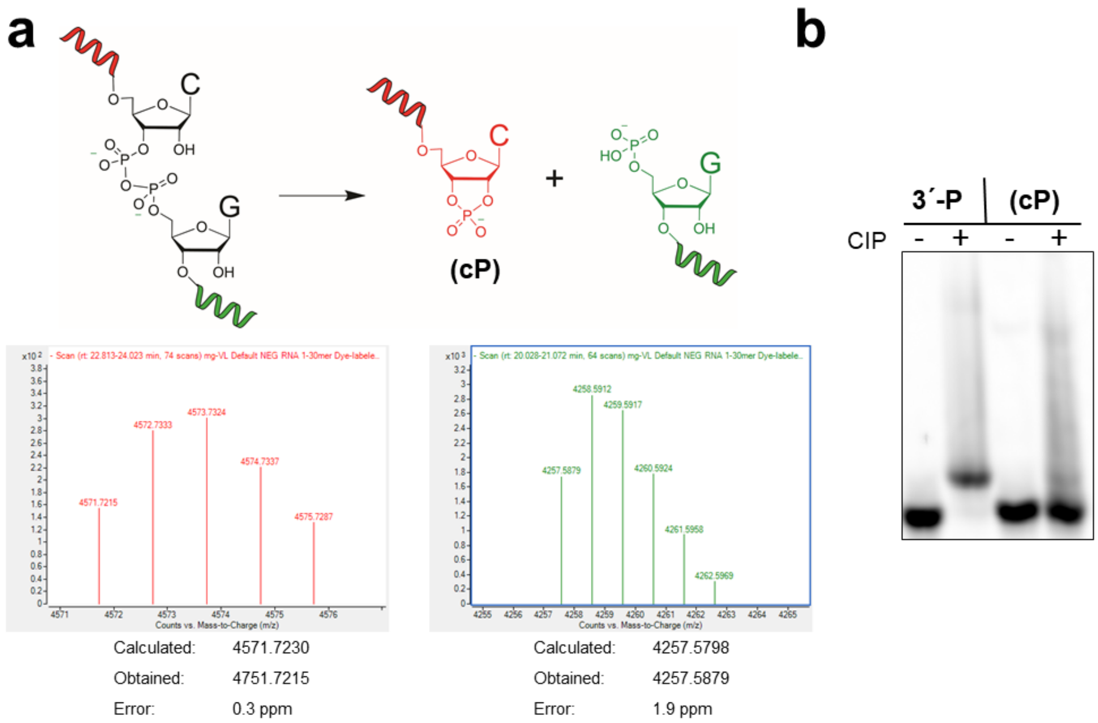

**Figure S2**. Characterization of the product of magnesium-catalyzed cleavage of ssRNA 25mer 1 containing a single pyrophosphate linkage. (a) Deconvoluted LC-MS mass spectrum of the two products of strand cleavage, a 5′ derived fragment bearing a terminal cyclic phosphate and a 3′ derived fragment containing a 5′-phosphate monoester. (b) Calf intestinal phosphatase (CIP) digestion of the cleaved product (5′ derived fragment) confirms the presence of a cyclic phosphate. The original 3′-phosphorylated primer (3′-P) is cleanly dephosphorylated by CIP treatment, whereas the cleaved product is unaffected; cyclic phosphate terminated RNA (cP) is not a substrate for CIP dephosphorylation.

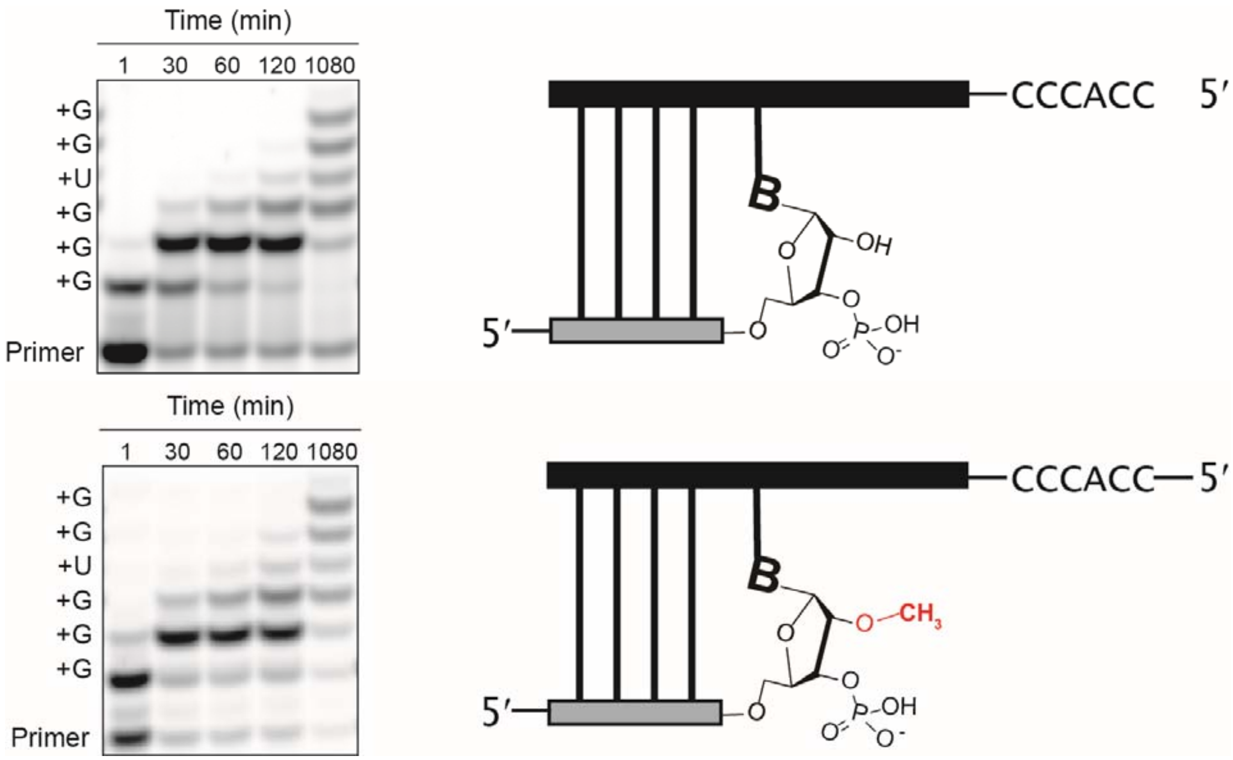

**Figure S3**. 2′-OMe modification of a 3′-phosphorylated primer eliminates cleavage of the pyrophosphate linkage during primer extension. The time course of primer extension with guanosine 2-aminophosphorimidazolide (2AImpG) and uridine-2-aminophosphorimidazolide (2AImpU) monomers was monitored using polyacrylamide gel electrophoresis (PAGE). The cleavage band is observed in the reaction with the 3′-phosphorylated primer (Top) as a band that comigrates with the original primer but does not change during the course of the assay. The reaction with 2′-OMe modification of the original primer displays the normal time-dependent loss of the primer band. All reactions were performed at pH 8.0, 200 mM HEPES, 50 mM $Mg^{2+}$ with 10 mM each guanosine 2-aminophosphorimidazolide and uridine-2-aminophosphorimidazolide.

#### 3. Supplementary tables

**Table S1. Magnesium dependence of the half-life of 3′-5′ pyrophosphate-linked RNA***

| $[Mg^{2+}]$ (mM) | 1 | 4 | 10 | 20 | 50 |
|---|---|---|---|---|---|
| $t_{1/2}$ (min) | 71(1) | 39(1) | 24(1) | 16(1) | 7.4(2) |

*The standard error of the mean is reported.

**Table S2. Sequences of oligonucleotides used in this study***

| Name | Sequence (5′→3′) | Comment |
|---|---|---|
| p-01 | /FAM/ AAG GGA AGA AGC 3p | 3′ phosphate primer |
| p-02 | /FAM/ AAG GGA AGA AGC 2p | 2′ phosphate primer |
| p-03 | /FAM/ AAG GGA AGA AGC | RNA primer |
| 2-01 | UAA UCC ACC CGC UUC UUC CCU U | Figures 2 and 3 template |
| 4-01 | TTG TCA TGC TAG CCC GCU TCT TCC CTT AAA A | Figure 4 template |
| 4-02 | TTG TCA TGC TAG ACC GCU TCT TCC CTT AAA A | Figure 4 template |
| 4-03 | 5p GCU AGC AUG AC | Figure 4 ligator |
| (1) | /FAM/ AAG GGA AGA AGC‡GGG CUA GCA UGA C | Pyrophosphate-linked template |
| (2) | /FAM/ AAG GGA AGA AGC‡G | Pyrophosphate-linked primer |
| (3) | /Cy3/ AUC GAA GGG‡GGC AAC ACG AC | Pyrophosphate-linked template |
| 5-01 | GGC GCU UCU UCC CUU AAA A | Figure 5 template |
| 5-02 | /FAM/ AAG GGA AGA AGC G | Figure 5 RNA primer |
| 5-03 | /FAM/ AAG GGA AGA AGC G 3p | Figure 5 3′ phosphate primer |
| 6-01 | /FAM/ GUC GUG UUG C | Figure 6a primer |
| 6-02 | /FAM/ GUC GUG UUG CC | Figure 6b primer |
| 6-03 | /FAM/ GUC GUG UUG CCC | Figure 6c primer |
| 6-04 | /Cy3/ AUC GAA GGG GGC AAC ACG AC | Figure 6 RNA template |
| S3-01 | /FAM/ AAG GGA AGA AGC 3p 2′OMe | Figure S3 2′OMe primer |

*Cy3 stands for Cyanine 3 dye; 5p stands for a terminal 5′-phosphate group; 3p stands for a terminal 3′-phosphate group; 2p stands for a terminal 2′-phosphate group; FAM stands for 6-fluorescein. ‡ symbolizes a 3′-5′ pyrophosphate linkage. The letters in red correspond to DNA nucleotides.
